# Regioselectively Acetylated Silybin Derivatives and their Cytotoxic Activity in Liver Cancer Cells

**DOI:** 10.1101/2025.02.13.638106

**Authors:** Tran Van Chien, Tran Thi Phuong Thao, Nguyen The Anh, Tran Van Loc, Michelle D. Garrett, Nguyen Thi Nga, Do Thi Thao, Christopher J. Serpell, Tran Van Sung

## Abstract

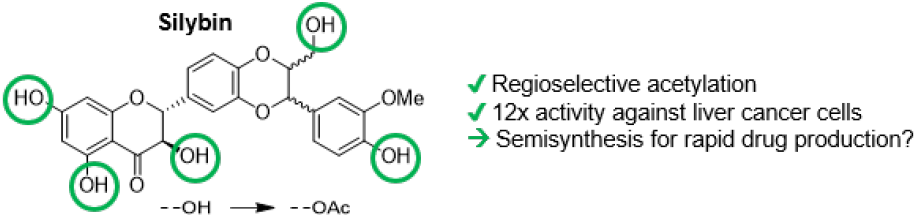

Natural products continue to serve as a valuable source of starting points for therapeutic agents. Six silybin derivatives have been synthesized and tested *in vitro* for their cytotoxic effect on the human hepatocellular carcinoma line HepG2 using the MTT assay. Results indicate that the 3,5,7,20,23-penta-*O*-acetyl-2,3-dehydrosilybin possesses a stronger cytotoxic effect than that of the parent silybin by a factor of 12. This compound was further evaluated for its cytotoxic activity on Hep3B, Huh7, and Huh7R hepatocellular carcinoma cell lines, giving similar potencies, arresting the cell cycle at S phase and causing apoptosis. These results illustrate the potential of silybin derivatives as therapeutics against liver cancer.

## Introduction

Silybin (also known as silibinin), one of the main flavonolignans of the herb *Silybum marianum*, has been used for the treatment of chronic liver diseases, including hepatitis, cirrhosis, and gallbladder disorders [1] [2], preventing liver cells from damage by chemicals or toxins by its high capacity for scavenging hydroxyl radicals [3] [4] [5]. Silybin also shows antifibrotic, anti-oxidant and anti-inflammatory activity, and inhibits hepatitis C virus RNA-dependent RNA polymerase activity [6] [7]. Recently, much attention has been paid to silybin due to its cytotoxic activity in various cancer cell lines, including small and non-small human lung carcinoma cells [8] [9], breast [10], bladder [11], prostate [12] [13], and human pancreatic cancers [14] [15]. Mechanistic experiments have suggested multifaceted routes of action, such as disruption of the cell cycle, decreasing the levels of cyclins (D1, D3, CDK2) [16] [17] [18] [19], inhibiting anti-apoptosis proteins (Bcl-2) and also growth factors (Flt, VEGF and IGF), or inhibiting angiogenesis and metastatic spread [20] [21] [22]. Recent studies on silybin have shown it can inhibit hepatocarcinogenesis and hepatocellular carcinoma growth by regulating the HGF/c-Met, Wnt/β-catenin, and PI3K/Akt/mTOR signalling pathways [23] [24] [25]. Silybin and its derivatives may therefore be promising agents for cancer therapy. However, the bioavailability and therapeutic efficiency of silybin itself are limited because of its poor water-solubility, and for this reason chemical modifications of silybin with the aim to improve the water-solubility but still remain its biological activity have been pursued.

In nature, silybin exists as an equal mixture of two diastereomers, silybin A and silybin B, which are inseparable by chromatographic methods. The presence of phenolic and alcoholic hydroxyl groups in silybin could serve as chemical handles for structural manipulations. Chemical modification and pharmacokinetic research on silybin derivatives have shown that acylation or alkylation of these hydroxyl groups may enhance the solubility of silybin, and help overcome the pharmacokinetic limitations of the parent compound [26].

As part of our ongoing efforts to develop naturally bioactive compounds for the treatment of liver cancer [27] [28] [29] [30], silybin was chosen as a promising candidate. To date, chemical modification on silybin has focused only on the hydroxyl groups on C-7 and C-23 [31] [32] [33] [34] [35], followed by evaluating their antioxidant potential. Only a small number of silybin derivatives have been synthesized and evaluated in cancer cells, especially in liver cancer cell lines in order to examine their possibility of treating liver-related diseases [33] [36] [37] [38]. Here, we report the synthesis of silybin derivatives by selective acetylation of hydroxyl groups across the molecule, and the evaluation of their biological activities in the HepG2 human hepatocarcinoma cell line. The most active compound was further investigated for its activity in Hep3B, Huh7, and Huh7R liver cancer cell lines, as well as its effects upon the cell cycle.

## Results and discussion

### Chemistry

Derivatization of silybin **1** (as a mixture of diastereomers, silybin A and silybin B) was initially carried out by the direct treatment with acetic anhydride in pyridine for 48 h at room temperature. After purification by silica gel column chromatography, the desired product **2** was obtained in 36% yield, together with **3** as a minor dehydrogenated product (18%) (Scheme 1). The structures of the products were determined based on the analysis of their NMR and MS data [26]. Compound **3** was assigned as 3,5,7,20,23-penta*-O-*acetyl-2,3-dehydrosilybin based on the absence of signals for protons H-2 and H-3 in its ^1^H NMR spectrum (appearing at *δ* 5.37/5.35 ppm (d, *J =* 12.0 Hz) and *δ* 5.69/5.67 ppm (d, *J =* 11.5 Hz for **2**) and the replacement of peaks at 81.0 ppm (C-2), and 73.4/73.2 ppm (C-3) in the ^13^C NMR spectrum of **2** with new olefin carbon signals at 154.8/154.1 ppm (C-2) and 133.5 ppm (C-3).

To synthesize silybin derivatives containing a free hydroxyl group at position 23 (23-OH), compound **1** was first converted into 23-*tert*-butyldimethylsilylsilybin by the treatment with *tert*-butyldimethylsilyl chloride (TBSCl) in pyridine at 50 °C, in the presence of AgNO_3_ as a catalyst. Subsequent acetylation, and removal of the TBS-protecting group provided products 3,5,7,20-tetra*-O-*acetylsilybin **4a** and 3,7,20-tri-*O-*acetylsilybin **4b**, in overall yields of 25% and 5%, respectively, after a simple chromatographic purification. Notably, removal of TBS-protecting group with either tetrabutylammonium fluoride (TBAF), or under acidic conditions (AcOH or HCOOH) in THF provided **4b** as a minor product, featuring removal of the acetyl group attached to 5-OH on silybin compared to solely **4a** which is the anticipated product based on published data [26] [39].

In second strategy for achieving new acetylation patterns, partially regioselective acetylation of silybin was achieved by exposure of silybin to acetic acid under Steglich esterification using the dehydrating reagent *N,N*′-dicyclohexylcarbodiimide (DCC) and 4-dimethylaminopyridine (DMAP) as a catalyst, yielding 3*-O-*acetylsilybin **5a** (40%) and 3,20-*di-O-*acetylsilybin **5b** (22%). This result is in contrast to reported esterification of silybin with carboxylic acids under Mitsunobu conditions, which forms the dehydrated product-hydnocarpin D [40], or the use of acyl chlorides in triethylamine (TEA) in tetrahydrofuran (THF), which esterifies preferentially the silybin 7-OH [38]. In our synthesis, the use of DCC/DMAP gave predominantly esterification of 3-OH of silybin, based on NMR analysis of the reaction products: in addition to proton and carbon signals similar to the parent silybin, the ^1^H NMR spectrum of **5a** contained a singlet peak corresponding to a methyl group at *δ* 2.08/2.07 ppm, together with peaks in the ^31^C spectrum corresponding to carboxylic ester and methyl signals at *δ* 169.5 ppm (CO) and 20.4 ppm (CO<u>C</u>H_3_), respectively. This confirms mono-acetylation of silybin. The ^1^H NMR spectrum also showed a downfield movement of the H-3 proton peak from *δ* 4.60 ppm in silybin (**1**) [41] to *δ* 5.75 ppm (t, *J =* 9.5 Hz, H-3), indicating the attachment of the acetyl group to 3-OH in silybin [26]. When compared with **5a**, it was clear that the NMR spectra of **5b** arose due to the addition of another acetyl group. We assigned the location of this second acetyl group to 20-OH of silybin, due to the downfield movement of the H-3 NMR peak at *δ* 5.75/5.74 ppm (d, *J* = 10.0 Hz) and the unchanged chemical shifts of H-6 [5.95/5.94 ppm (d, *J* = 2.0 Hz, H-6)] and H-8 [6.0 ppm (d, *J* = 2.0 Hz, H-8)] compared to silybin [41], along with the clear appearance of carbon signals at *d*_C_ 151.6 ppm (C-19) [downfield from *δ* 147 ppm in silybin] and 140.3 (C-20) [upfield from *δ* 146.5 ppm in silybin], indicating that the secondary acetyl group attached to 20-OH [26].

**Scheme 1.**
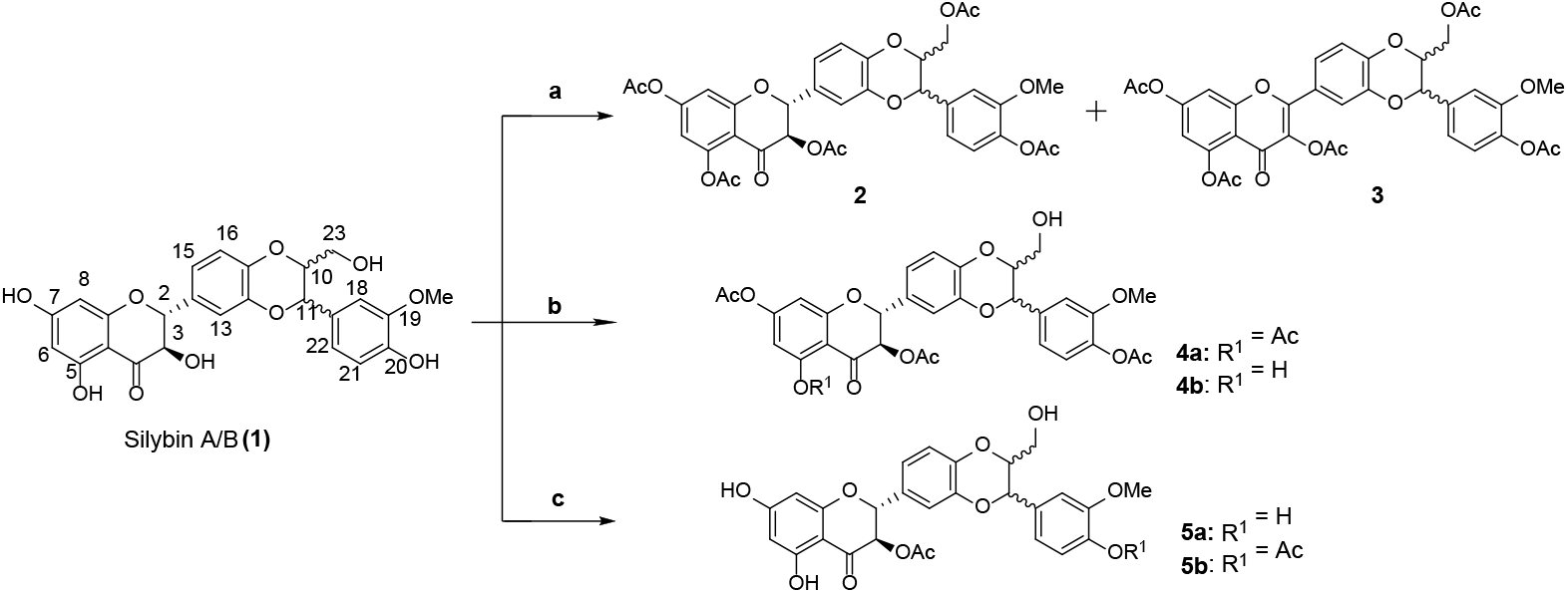
Synthesis of silybin derivatives. Reagents and conditions: a) Ac_2_O, pyridine, rt, 48 h, 36% **(2)**, 18% **(3)**, b) i) TBSCl, AgNO_3_, Pyridine, 50 oC, 4 h; ii) Ac_2_O, pyridine, rt, 14 h; iii) AcOH, THF/H_2_O (1/1), 25% (**4a**, three steps), 5% (**4b**, three steps), c) AcOH, DCC, DMAP, THF, rt, 12 h, 40% (**5a**) and 22% (**5b**).

### Bioactivity

#### Cytotoxic activity

Silybin and its derivatives were tested *in vitro* for their cytotoxic activity on the HepG2 human hepatocellular (liver) carcinoma cell line, with ellipticine used as a positive control. As presented in Table 1, all silybin derivatives, except for compound **5a**, exhibited considerably stronger cytotoxic potency on HepG2 cells than that of silybin. While there is a general trend of higher acetylation levels giving lower IC_50_ values, there are considerable fluctuations. Compound **2** (with complete acetylation of the hydroxyl groups) showed cytotoxicity 4-fold stronger than that of silybin, while partial acetylation exemplified by **4a** and **4b** enhanced the cytotoxic potency by approximately 2- and 3-fold on HepG2 compared to silybin (**1**). The double acetylated **5b**, demonstrates 3-fold greater cytotoxicity compared to monoacetylated **5a**.

**Table 1.**
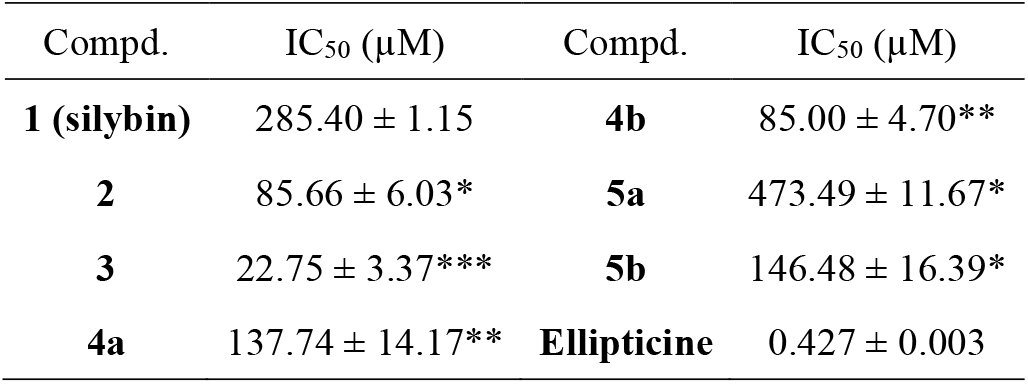
Cytotoxic activity of silybin (**1**) and its derivatives (**2-5**) on the HepG2 human hepatocellular (liver) carcinoma cell line (IC_50_ (µM), 96 hour MTT assay, n ≥ 3 assay repeats), *p<0.05; **p<0.01; ***p<0.001

However, there was an opposite cytotoxic relationship for **4a** and **4b** in relation to acetylation number, where the monoacetylated **5a** was less cytotoxic on HepG2 cells than silybin. The fact that the location of the modification matters suggests that the effects are due to more than physicochemical modulation of the compound. Of all the compounds, 3,5,7,20,23-penta-*O*-acetyl-2,3-dehydrosilybin (**3**) exhibited the most potent inhibition of HepG2 cancer cell proliferation with an IC_50_ value of 22.75 µM, which was approximately 3.7-fold more potent than that of compound **2**; and 12.5-fold better cytotoxic potency compared to silybin (**1**), indicating that the presence of a 2,3-double bond in 2,3-dehydrosilybin derivatives significantly increases the cytotoxicity towards HepG2 cells.

We selected 3,5,7,20,23-penta-*O*-acetyl-2,3-dehydrosilybin (**3**) for evaluation against three further human hepatocellular carcinoma cell lines (Hep3B, Huh7, and Huh7R) because of its high cytotoxicity on HepG2 cells, again using the MTT assay (Fig. 1). The cytotoxicity of **3** was significantly more potent on Huh7 (IC_50_ = 13.67 µM) when compared with other hepatic carcinoma cell lines, Hep3B and Huh7R cells, with the IC_50_ values of 17.09 (p<0.05) and 18.59 µM (p<0.01), respectively.

**Figure 1.**
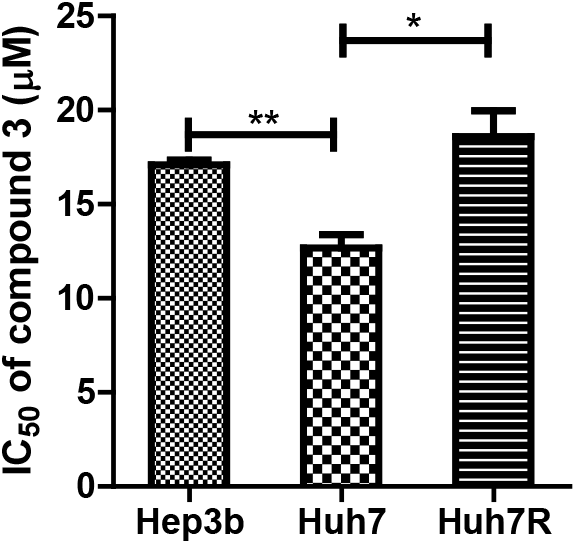
The antiproliferation capacity of 3,5,7,20,23-penta-*O*-acetyl-2,3-dehydrosilybin (**3**) on Hep3B, Huh7 and Hub7R hepatocellular carcinoma cell lines. *p<0.05; **p<0.01

### Cell cycle analysis

In order to gain insights into the mechanism of action behind the antiproliferative effects of 3,5,7,20,23-penta-*O*-acetyl-2,3-dehydrosilybin (**3**), HepG2 cells were treated with the compound at 1 x IC_50_ and 3 x IC_50_ µM for 24 h, and the cell cycle distribution was then measured by flow cytometry. The results indicated clearly that compound **3** caused a substantial arrest of the cell cycle in HepG2 cells at S phase in a dose-dependent manner. A total of 62.73% of cells were observed at S phase at 3 x IC_50_, while 29.55% were in the same phase at 1 x IC_50_, and just 20.20 % in the negative control (Figure 2).

**Figure 2.**
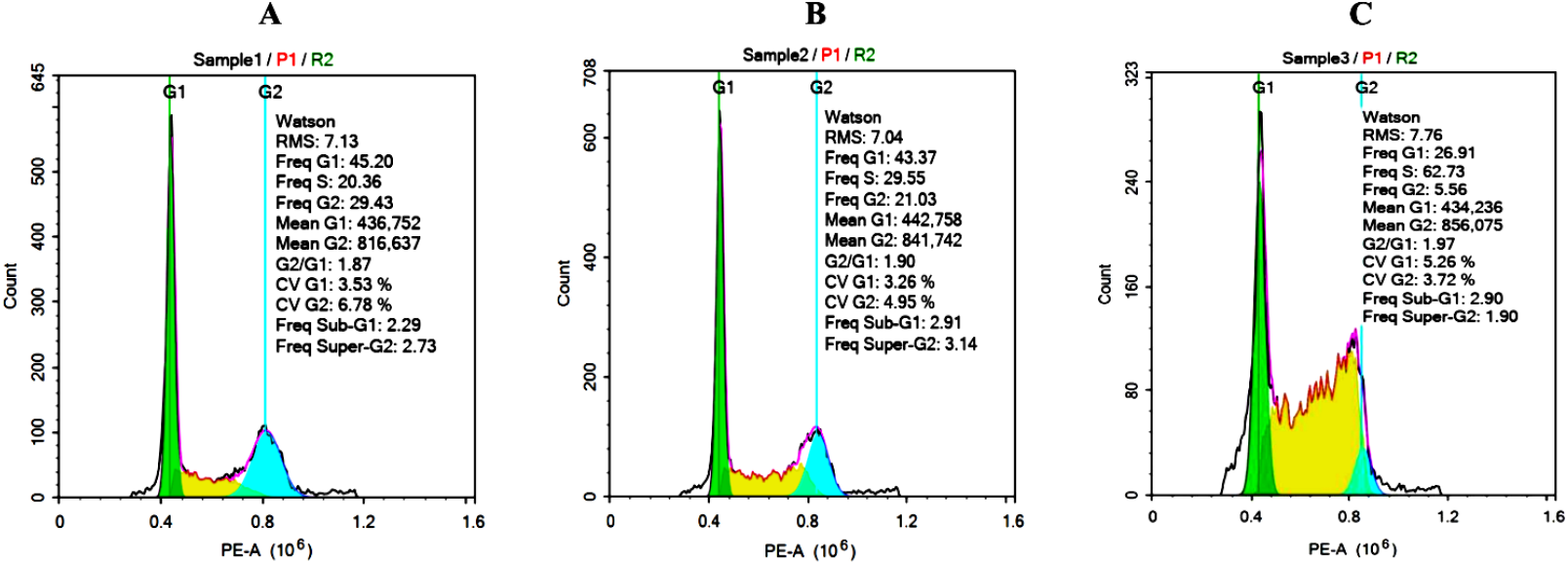
The effect of **3** on the cell cycle profile of HepG2 cells. HepG2 cells were incubated for 24 hours with (A) cell cultured media alone (negative control); (B) with (**3**) at 1 x IC_50_; (C) **3** at 3 x IC_50_ followed by cell cycle analysis.

### Apoptosis studies

Apoptosis is a critical pathway that leads to cancer cell death and so the extent of apoptosis of HepG2 cells was determined via Annexin V staining after incubation with compound **3** for 24 hours. It can be seen that **3** induces apoptosis in a dose-dependent manner (Figure 3, quadrant Q2-2). Specifically, there was in an approximately two-fold increase in the number of apoptosis cells in the group treated with **3** at 3 x IC_50_ compared to 1 x IC_50_, and accordingly, the number of cells that enter apoptosis in the 1 x IC_50_ group was higher than that in the negative control. However, **3** not only induces cell death by apoptosis, but also causes necrosis. At a concentration of 3 x IC_50_, **3** notably increased the percentage of necrotic cells (quadrant Q2-1) when compared with negative control and the lower concentration treatment.

**Figure 3.**
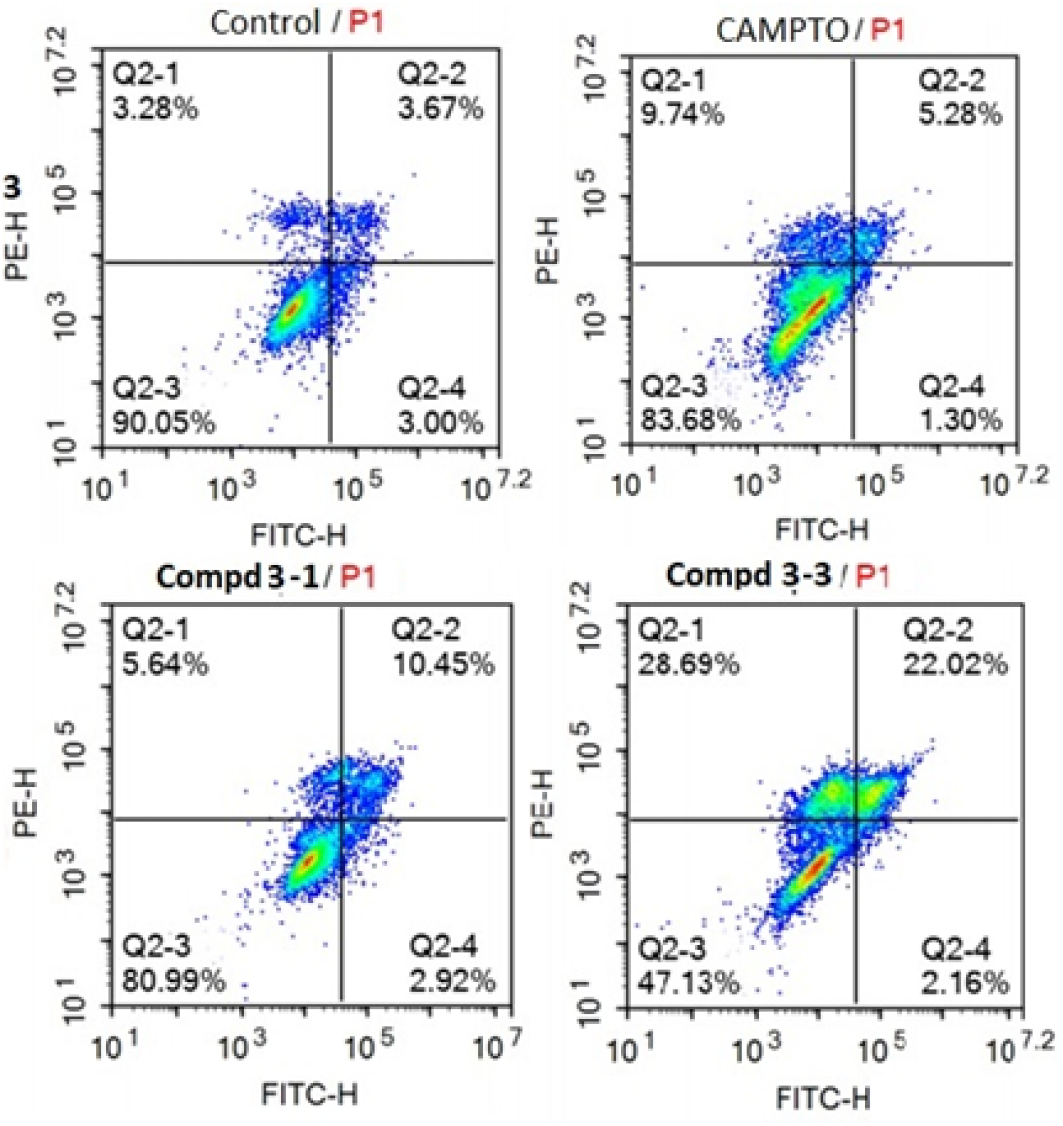
: The effect of **3** on induction of apoptosis in HepG2 cells. HepG2 cells were incubated for 24 hours with cell culture media alone (negative control), 0.5 μM Camptothecin (CAMPTO, positive control), or 3,5,7,20,23-penta-*O*-acetyl-2,3-dehydrosilybin (**3**) (**3**-**1** indicating compound **3** at concentration of 1 x IC_50_; and **3**-**3** indicating compound **3** at concentration of 3 x IC_50_) followed by detection of Annexin V staining via flow cytometry.

Measurement of caspase 3 activation is an alternative way to evaluate compound-induced apoptosis and so the effect of **3** on caspase activity was measured in HepG2 cells after 1, 6 and 24 hours of treatment. It can be seen that caspase 3 activity in HepG2 was similar to the control after 1 hour of treatment with concentration at 0.3 x IC_50_ and 1 x IC_50_ (µM) and declined at 6 and 24 hours (Table 3). However, the activity of caspase 3 increased after treatment with concentration at 3 x IC_50_ (µM) of **3** at all three time points. *p<0.05; **p<0.01 compared with control.

## Discussion

It was observed that five of the six silybin derivatives demonstrated greater cytotoxic activity in the HepG2 human hepatocellular carcinoma cell line in comparison to silybin. Among the tested derivatives, 3,5,7,20,23-penta-*O*-acetyl-2,3-dehydrosilybin (**3**) showed the highest cytotoxicity on the HepG2 cell line (IC_50_: 22.75 µM), and exhibited similar cytotoxicity to this on additional human hepatocellular carcinoma cell lines Hep3B, Huh7, and Huh7R cell lines (Table 2 and Fig 1). Compound **3** also induced an S - phase arrest in HepG2 cells. In contrast Varghese and colleagues reported that silybin induced a G1 arrest at 24 hours post-treatment suggesting that the mechanism of action of **3** may be distinct from silybin. The distinct stages of the cell cycle (G1, S, G2, and M) play an important role in the normal survival of cells, and the initiation of each stage relies on cell cycle checkpoints, which ensure the proper proliferation [24]. This DNA surveillance mechanism will not allow cell division if the accumulation of genetic errors is not controlled, and leads to cell apoptosis in that case [42]. Compound **3** also led HepG2 to enter apoptosis. Specifically, compound **3** induced apoptosis in HepG2 cells at 1 x IC_50_, as determined by Annexin V staining which is much lower than the assay concentrations reported for silybin (100 - 300 µM) by Varghese and colleagues, who also used this method to measure apoptosis [17]. It has also been reported that the apoptotic rate of silybin on HepG2 cells was 60% at a concentration of 68 µM, but this was at 72 hours post treatment and so cannot be directly compared with results for **3** [43]. Apoptosis has been defined as a cell’s natural mechanism for programmed cell death. This pathway is executed through activated initiator and executioner caspases, resulting in DNA damage and cell death [44]. Therefore, targeting the apoptosis process is a potential approach for cancer therapies because it eliminates the uncontrolled growth of malignant cells. In our study, compound **3** also gave rise to an increase in caspase 3 activation. Caspase 3, together with caspases 6 and 7, is the executioner caspase or effector caspase, which is responsible for cleaving variation of downstream substrates such as inhibitor of caspase-activated DNAse (ICAD), Rho-associated coiled-coil protein kinase 1 (ROCK1). These proteins conduct to typical morphological changes in apoptotic cells [45] [46]. This activity once again confirms the apoptotic effect of compound **3** on the HepG2 human hepatocellular carcinoma cell line.

**Table 2.** Cytotoxic activity of 3,5,7,20,23-penta-*O*-acetyl-2,3-dehydrosilybin (**3**) on Hep3B, Huh7, and Huh7R human hepatocellular carcinoma cell lines (IC_50_ (µM), 96 hour MTT assay, n ≥ 3 assay repeats).

| Compd. | Hep3B | Huh7 | Huh7R |
| --- | --- | --- | --- |
| **3** | 17.09±0.26 | 12.67±0.70 | 18.59±1.38 |

**Table 3.** Caspase 3 induction in HepG2 cells treated with 3,5,7,20,23-penta-*O*-acetyl-2,3-dehydrosilybin (**3**)

| Compd. | Conc<br>(μM) | Fold change in caspase 3 activity * | | | | | |
| --- | --- | --- | --- | --- | --- | --- | --- |
|  |  | 1 h |  | 6 h |  | 24 h |  |
|  |  | Mean | SD | Mean | SD | Mean | SD |
| **3** | 0.3 x IC<sub>50</sub> | 1.08 | 0.17 | 0.73* | 0.07 | 0.63* | 0.07 |
|  | 1 x IC<sub>50</sub> | 1.18 | 0.09 | 0.77* | 0.06 | 0.74* | 0.05 |
|  | 3 x IC<sub>50</sub> | 1.47* | 0.10 | 1.29* | 0.09 | 1.26 | 0.12 |
| **Camptothecin** | 0.5 | 1.94** | 0.08 | 2.21** | 0.11 | 2.19** | 0.14 |
| **Control** |  | 1.00 | 0.08 | 1.00 | 0.05 | 1.00 | 0.02 |

\*Versus untreated control at each time point

## Conclusions

Six silybin derivatives (four of which are novel) have been synthesized and evaluated for their biological activity on the HepG2 human hepatocellular carcinoma cell line. Five of six compounds exhibited greater cytotoxic activity versus silybin itself. In particular, the 2,3-dehydrosilybin derivative **3** exhibited strong cytotoxic activity on HepG2, Hep3B, Huh7 and Huh7R human hepatocellular carcinoma cell lines. This compound caused a cell cycle arrest in HepG2 cells at the S phase in a dose-dependent manner. Compound **3** also induced apoptosis in the HepG2 cells as examined by Annexin V staining and caspase 3 activation. These results may provide valuable insights into the potential chemopreventive efficacy of 3,5,7,20,23-penta-*O*-acetyl-2,3-dehydrosilybin (**3**) in the HepG2 cell line. This study found that the anticancer activity of this compound was mediated through the decline of cell viability, inhibition of the cell cycle, and activation of apoptosis through the rising level of caspase 3. The further development of silybin derivatives, combined with deeper mechanistic studies may provide a route to new compounds which are potent against liver cancer.

## Acknowledgments

We thank the UK’s Engineering and Physical Science Research Council and Global Challenges Research Fund (EP/T020164/1) and the University of Kent, United Kingdom and Vietnam Academy of Science and Technology (VAST) for financial support under the project.

## 3. Experimental

Chemicals were purchased from Sigma-Aldrich and used without further purification. Solvents were redistilled before being used. NMR spectra (^1^H, ^13^C, HMBC, HSQC) were recorded on a Bruker AVANCE 500 MHz with tetramethylsilane (TMS) as the internal standard. Chemical shifts are reported in parts per million (δ ppm). *J* coupling constants in Hz. Proton spectra multiplicities are abbreviated as: s singlet, brs broad single, d doublet, t triplet, q quartet, quin quintet, m multiplet, dd double doublet, dt double of triplet, td triplet of double. Electrospray ionization (ESI) mass spectra were measured on a 1100 Agilent LC/MS ion trap. Reactions were monitored by thin-layer chromatography using silica gel G60 F254 (Merck). Silica gel 300–400 mesh (Merck) was used for column chromatography.

Hepato-carcinoma cell lines (HepG2, Hep3B, Huh and Huh7R) were the kindly gift of Prof. Chi-Ying Huang, Taiwan. DMEM (Gibco, USA) consisted of 10% fetal bovine serum (FBS, Gibco), 2mM L-Glutmine (Gibco, USA) and 1% Anti-Anti was used for growth cancer cells. Huh7R particularly was cultured in DMEM with Sonitanib (5µM). 3-(4,5-Dimethylthiazol-2-yl)-2,5-diphenyltetrazolium bromide (MTT), dimethyl sulfoxide (DMSO), ribonuclease (RNase), and propidium iodide (PI) were origin from Sigma Chemical (St. Louis, MO, USA). The eBioessence™ Annexin V Apoptosis Detection Kit was purchased from Invitrogen (Carlsbad, CA, USA). Caspase-3 assay kit (colorimetric) sourced from Abcam (Cambridge, UK).

### Synthesis of silybin derivatives 2 and 3

Silybin (mixture of silybin A and B, 1:1) (**1**, 240 mg, 0.5 mmol) was treated with acetic anhydride (2 mL) in pyridine (2 mL) at room temperature for 48 h. The reaction mixture was concentrated under reduced pressure to dryness, and followed by purification over a silica gel column chromatography (*n*-Hexane/EtOAc; 10/1) to yield product **2** (125 mg, 36%) and **3** (62 mg, 18%), respectively.

*3,5,7,20,23-Penta-O-acetylsilybin* ***(2)***: [26] [39] Yellow amorphous powder; *R*_*f*_ = 0.25 (*n*-Hexane/EtOAc; 10/1). ESI-MS *m/z*: 715.1 [M+Na]^+. 1^H-NMR (500 MHz, CDCl_3_) *δ* 7.12-7.08 (m, 2H, H-13, H-15), *δ* 7.02-6.96 (m, 4H, H-16, H-18, H-21 and H-22), 6.78/6.77(d, *J =* 2.5 Hz, H-8), 6.59/6.58 (d, *J =* 2.5 Hz, H-6), 5.69/5.67 (d, *J =* 12.0, H-3), 5.37/5.35 (d, *J =* 11.5, H-2), 4.95 (dd, *J =* 8.0, 2.0 Hz, H-11), 4.37 (dt, *J =* 12.5, 2.0, 1H, H-23), 4.28-4.25 (m, 1H, H-10), 4.02 (dt, *J =* 12.5, 2.0, 1H, H-23),3.86/3.85 (s, 3H, 19-OCH_3_), 2.37 (s, 3H, 5-OAc), 2.32 (s, 3H, 7-OAc), 2.28 (s, 3H, 20-OAc), 2.06 (s, 3H, 23-OAc), 2.05 (s, 3H, 3-OAc). ^13^C-NMR (125 MHz, CDCl_3_) *δ* 185.3 (C-3), 170.4, 169.2, 168.7, 167.8, 162.5, 156.4, 151.7, 151.5, 143.9, 143.6/143.5, 140.6, 134.4, 128.6/128.5, 123.2, 121.1, 120.9, 119.9, 117.4/117.3, 116.4/116.3, 111.2, 111.1, 110.6, 108.9, 81.0 (C-2), 76.4/76.3 (C-10), 75.6 (C-11), 73.4/73.2 (C-3), 62.6 (C-23), 56.1 (OMe), 21.2 (7-OAc), 20.9 (5-OAc), 20.7 (20-OAc), 20.6 (23-OAc), 20.4 (3-OAc).

*3,5,7,20,23-Penta-O-acetyl-2,3-dehydrosilybin* ***(3)***: [26] White amorphous powder; *R*_*f*_ = 0.40 (*n*-Hexane/EtOAc; 10/1). ESI-MS *m/z*: 713.2 [M+Na]^+. 1^H-NMR (500 MHz, CDCl_3_) *δ* 7.52 (d, *J =* 2.0 Hz, H-8), 7.46 (dd, *J =* 7.0, 1.5 Hz, H-15), 7.31 (d, *J =* 2.0 Hz, H-6), 7.11 (d, *J =* 6.5 Hz, H-22), 7.09 (d, *J =* 7.5 Hz, H-21), 7.00-6.98 (m, 2H, H-16, H-13), 6.85 (d, *J =* 2.0 Hz, H-18), 4.99 (d, *J =* 7.0 Hz, H-11), 4.41 (dd, *J =* 5.5, 3.0 Hz, 1H, H-23), 4.35 (brs, 1H, OH), 4.34-4.32 (m, 1H, H-10), 4.03 (dd, *J =* 5.5, 3.0 Hz, 1H, H-23), 3.87 (s, 3H, OMe), 2.43 (s, 3H, 3-OAc), 2.34 (s, 6H, 2xOAc), 2.32 (s, 3H, 20-OAc), 2.08 (s, 3H, 23-OAc). ^13^C-NMR (125 MHz, CDCl_3_) *δ* 179.6, 170.3, 170.2, 169.3, 168.7, 167.9, 167.8, 162.7, 156.9, 154.8, 154.1, 151.8, 150.4, 145.8, 143.5, 140.7, 134.1, 133.5, 123.4, 122.9, 122.4, 119.9, 117.5, 117.4, 114.7, 113.7, 111.1, 108.9, 76.4 (10-C), 75.9 (11-C), 62.5 (23-C), 56.0 (OMe), 21.2 (OAc), 21.1 (OAc), 20.7 (OAc), 20.6 (2xOAc).

### Synthesis of silybin derivatives 4a-b

Silybin (580 mg, 1.2 mmol) was dissolved at room temperature in dry pyridine (5 mL), and *tert-* butyldimethylsilyl chloride (TBDMSCl, 181 mg, 1.2 mmol) and AgNO_3_ (50 mg, 0.3 mmol) were added. After being stirred at 50 °C for 4 h, the reaction mixture was quenched with ethyl acetate (150 mL) and wahsed with brine solution (3 × 10 mL). The solvent was dried over Na_2_SO_4_ and evaporated under vacuum. A silica gel column chromatography (*n*-Hexane/EtOAc; 3/1) provided the intermediate product 23-*tert-*butyldimethylsilylsilybin (415 mg, 58%), as a white amorphous powder. The intermediate product (415 mg, 0.7 mmol) was subsequently exposed to acetic anhydride (1.5 mL) in pyridine (5 mL). The reaction mixture was stirred 24 h at room temperature, was then added a saturated solution of NaHCO_3_ (10 mL), and extracted with EtOAc (2 × 100 mL). The organic phase was combined and washed with brine solution, dried over Na_2_SO_4_ and evaporated to dryness. The residue was dissolved in the mixture of THF/H_2_O (6 mL, 1/1), AcOH 10% (0.5 mL) was added and the mixture was stirred further 6 h at room temperature. Afterwards, the reaction mixture was diluted with EtOAc (150 mL), washed with water, dried over Na_2_SO_4_. Removal of the solvent and purification over a silica gel column chromatography (*n*-Hexane/EtOAc; 5:1) yielded compound **4a** (195 mg, 25%) and **4b** (28 mg, 4%) over three-step.

*3,5,7,20-Tetra-O-acetylsilybin* ***(4a)***: Yellow amorphous powder; *R*_*f*_ = 0.30 (*n*-Hexane/EtOAc; 6/1). ESI-MS *m/z*: 673.0 [M+Na]^+^_. 1_H-NMR (500 MHz, CDCl_3_) *δ* 7.12-6.97 (m, 6H, H-13, H-15, H-16, H-18, H-21 and H-22), 6.79/6.77 (d, *J =* 2.5 Hz, H-8), 6.59/6.58 (d, *J =* 2.5 Hz, H-6), 5.69 (t, *J =* 12.0 Hz, H-3), 5.37/5.35 (d, *J =* 12.5 Hz, H-2), 5.05 (dd, *J =* 8.0, 2.0 Hz, H-11), 4.06-4.03 (m, 1H, H-10), 3.87/3.86 (s, 3H, 19-OCH_3_), 3.85 (1H, overlap, H-23), 3.60 (dd, *J =* 13.0, 4.0 Hz, 1H, H-23), 2.37 (s, 3H, 5-OAc), 2.33 (s, 3H, 20-OAc), 2.29 (s, 3H, 7-OAc), 2.20-2.12 (m, 1H, 23-OH), 2.05 (s, 3H, 3-OAc). ^13^C-NMR (125 MHz, CDCl_3_) *δ* 185.3 (C-4), 171.3, 169.2, 168.9, 167.9, 162.6/162.5 (C-5), 156.4 (C-7), 151.6/151.5 (C-8a or C-19), 144.3 (C-12a), 143.8 (C-16a), 140.4 (C-20), 134.7 (C-17), 128.5 (C-14), 123.2/123.1 (C-21), 121.0 (C-15), 119.9/119.8 (C-22), 117.3/117.2 (C-16), 116.4 (C-13), 111.4 (C-18), 111.2 (C-8), 110.6 (C-4a), 109.0 (C-6), 81.1/81.0 (C-2), 78.4/78.3 (C-10), 76.0 (C-11), 73.4/73.2 (C-3), 61.5 (C-23), 56.1 (19-OCH_3_), 21.2 (7-OAc), 21.0 (5-OAc), 20.7 (20-OAc), 20.5 (3-OAc).

*3,7,20-Tri-O-acetylsilybin* ***(4b)***: [26] Yellow amorphous powder; *R*_*f*_ = 0.20 (*n*-Hexane/EtOAc; 6/1). ESI-MS *m/z*: 607.0 [M-H]^-^_. 1_H-NMR (500 MHz, CD_3_OD) *δ* 7.21 (1H, m, H-21), 7.24-7.03 (6H, H-20, H-13, H-15, H-16, H-18 and H-22), 6.36 (d, *J =* 2.0 Hz, H-8), 6.26 (d, *J =* 2.0 Hz, H-6), 5.70/5.68 (d, *J =* 12.0, H-3), 5.35 (d, *J =* 12.0, H-2), 5.07 (d, *J =* 8.0 Hz, H-11), 4.15-4.13 (m, 1H, H-10), 3.86/3.85 (3H, s, 19-OCH_3_), 3.94 (dd, *J* = 12.0, 2.5 Hz, 1H, H-23), 3.65 (dd, *J =* 12.0, 2.5 Hz, 1H, H-23), 2.32 (s, 3H, 7-OAc), 2.29 (s, 3H, 20-OAc), 2.00 (s, 3H, 3-OAc).

### Synthesis of silybin derivatives 5a-b

A stirred solution of silybin (120 mg, 0.25 mmol) and AcOH (20 mg, 0.3 mmol) in THF (2 mL) at 0 °C was added *N,N*′-dicyclohexylcarbodiimide (DCC, 75 mg, 0.36 mmol) and 4-dimethylaminopyridine (DMAP, 36 mg, 0.3 mmol). The reaction mixture was stirred at room temperature for 48 h, then an amount of oxalic acid (0.3 equiv.) was added slowly to scavenge the excess amount of DCC. The reaction was stirred for 15 min and cooled to -5-0 °C for 1 h. The precipitated solid was removed by filtration. Removal of solvent and purification over a silica gel column chromatography, eluting with DCM/Acetone (5/1) yielded compound **5a** (52 mg, 40%) and **5b** (31 mg, 22%), respectively.

*3-O-Acetylsilybin* ***(5a)***: White amorphous powder; *R*_*f*_ = 0.22 (DCM/Acetone; 5/1). ESI-MS *m/z*: 525.0 [M+H] ^+. 1^H-NMR (600 MHz, CDCl_3_) *δ* 11.47 (s, 1H, 5-OH), 7.13-7.11 (m, 1H, H-13), 7.11-6.92 (m, 5H), 6.48 (brs, 1H, OH), 6.04/6.03 (s, 1H, H-8), 5.98/5.97 (s, 1H, H-6), 5.75 (t, *J =* 12.0 Hz, H-3), 5.24 (dd, *J =* 12.0, 5.4 Hz, H-2), 4.95 (d, *J =* 8.4 Hz, H-11), 4.07-4.05 (m, H-10), 3.93/3.92 (s, 3H, 19-OCH_3_), 3.81 (d, *J =* 12.6 Hz, 1H, H-23), 3.57 (dd, *J =* 12.6, 4.2 Hz, 1H, H-23), 2.08/2.07 (s, 3H, 3-OAc). ^13^C-NMR (150 MHz, CDCl_3_) *δ* 191.6 (C-4), 169.5/169.4 (OAc), 165.4 (C-7), 164.3 (C-5), 162 (C-8a), 147.0 (C-19), 146.6 (C-20), 144.2 (C-16a), 143.9 (C-12a), 128.7 (C-14), 127.7 (C-17), 120.8 (C-15), 120.6 (C-22), 117.2/117.1 (C-16), 116.4 (C-13), 114.7/114.6 (C-21), 109.6/109.5 (C-18), 100.6/109.5 (C-4a), 101.9, 97.3 (C-6), 95.8 (C-8), 80.9 (C-2), 78.3 (C-10), 76.4 (C-11), 72.5/72.4 (C-3), 61.7 (C-23), 56.1 (OCH_3_), 20.4 (3-OAc).

*3,20-Di-O-acetylsilybin* ***(5b)***: White amorphous powder; *R*_*f*_ = 0.32 (DCM/Acetone; 5/1). ESI-MS *m/z*: 567.0 [M+H] ^+. 1^H-NMR (600 MHz, CDCl_3_) *δ* 11.46 (s, 1H, 5-OH), 7.13-7.11 (m, 1H, H-13), 7.09-6.95 (m, 5H), 6.88 (brs, 1H, OH), 6.02/6.01 (s, H-8), 5.96/5.95 (s, H-6), 5.75 (t, *J =* 14.4 Hz, H-3), 5.27 (dd, *J =* 12.0, 5.4 Hz, H-2), 5.03 (d, *J =* 8.4 Hz, H-11), 4.06-4.02 (m, 1H, H-10), 3.86/3.85 (s, 3H, 19-OCH_3_), 3.85 (1H, overlap, H-23), 3.58 (m, 1H, H-23), 2.33 (s, 3H, 20-OAc), 2.08/2.07 (s, 3H, 3-OAc). ^13^C-NMR (150 MHz, CDCl_3_) *δ* 191.5, 169.5, 169.2, 165.9/165.7, 162.9/162.4, 151.5, 146.6/146.5, 144.1, 143.8, 140.3, 134.8, 128.8/128.7, 123.1, 120.9, 119.8, 117.2, 116.3, 114.8, 111.3, 101.7, 97.7, 95.9, 80.8, 78.3/78.2, 75.9, 72.5, 61.5, 56.1, 20.7 (20-OAc), 20.4 (3-OAc).

### Bioassay

#### Growth inhibition assay

Liver cancer cell lines HepG2, Hep3B, Huh7 and Huh7R were grown in DMEM containing 10% FBS and 1% antibiotic (Anti-Anti, Gibco, Thermo Fisher Scienctific) in a humidified atmosphere at 37 °C and 5% CO_2_. In order to test the cytotoxic effect of compounds on different cell lines, cells were detached from the adherent culture surface by Trypsin – EDTA (0.05%) and then seeded into 96 wells-plate at the density of 3×10^4^ cell/mL and treated with an eight titration of each compound. Ellipticine was used as a reference (positive) control. After 96 hours of incubation, all cell culture media was removed from wells followed by addition of 100 µL MTT solution (0.5 mg/mL solute in fresh medium) per well and incubation at 37 °C for 4 hours. The MTT solution was then removed and DMSO (200 µL) to each well to solubilise the formazan product and mix each sample again using a pipette before determining the absorbance at 570 nm. GI_50_ values (compound concentration that reduces the MTT assay value by 50% versus untreated control) were calculated using GraphPad Prism 5.0 software.

#### Cell cycle determination

To evaluate cell cycle effects, HepG2 cells were seeded in T25 flasks at a density of 1×10^5^ cell/mL at 37 °C in 5% CO_2_ and treated with compound at concentrations of 1 x IC_50_ and 3 x IC_50_ from the result of the cytotoxic assay. After 24 hours of treatment, cells were gently harvested with trypsin and then washed with PBS (pH 7,4). In the next step, 70% ethanol was slowly added to the cells after which they were kept in a fridge (4 °C) for at least 2h to fix the cells. After fixation the cells were pelleted by centrifugation and the ethanol completely removed resuspension with RNase A (1 mg/mL) and incubation at 37 °C for 15 min, followed by PI staining for 45 min. Finally, at least 10,000 cells were analysed using a NovoCyte flow cytometer system with NovoExpress software (ACEA Bioscience Inc.) to generate cell cycle profiles.

#### Annexin V staining for apoptosis

The eBioessenceTM Annexin V Apoptosis Detection Kit was used to measure the percentage of apoptotic cells after treatment with each compound for 24 hours according to the manufacturer’s instructions. Briefly, after harvesting by trypsin-EDTA, cells were washed with cold PBS to completely remove trace medium. Cell pellets were resuspended in binding buffer and then stained with Annexin V-FITC for 15 minutes at room temperature (protected from light) before being washed with the binding buffer to remove the unstained dye, resuspension in 190 µL of binding buffer and addition of 10 µL of PI solution (20 µg/mL) in a binding buffer. Analysis was performed using a NovoCyte flow cytometer system and NovoExpress software (ACEA Bioscience Inc.) to identify the proportion of apoptotic cells.

#### Caspase 3 inducible assay

A Caspase-3 Colorimetric Assay Kit (Biovision Inc.) was used for establishing the caspase 3 activity of tested samples. Treated cells were lysed in lysis buffer for 10 minutes in ice and centrifuged at 10,000 x g for 1 minute to remove residual cellular debris (cell pellet). After determination of the protein concentration of each sample (Bradford assay), 80 µg of each sample in 50 µL assay buffer was added to 50 µL DTT (10 mM) and 5 µL of DEVD-pNA (200 μM) in each well of a 96-well plate in triplicate. The plate was incubated at 37 °C for 1 hour. The absorbance was read at 405 nm using a microplate reader (BioTek, ELx800).

#### Statistical analysis

Results were analysed using Excel and GraphPad Prism 5.0 software and reported as mean ± standard error (SE). GraphPad Prism 5 software using unpaired t-test and one-way analysis of variance was used for data analysis. A value of *P* < 0.05 was considered to indicate statistical significance.

